# What are the reference strains of *Acinetobacter baumannii* referring to?

**DOI:** 10.1101/2022.02.27.482139

**Authors:** Chantal Philippe, Adam Valcek, Clémence Whiteway, Etienne Robino, Kristina Nesporova, Mona Bové, Tom Coenye, Tim De Pooter, Wouter De Coster, Mojca Strazisar, Johanna Kenyon, Charles Van der Henst

**Author notes:** These authors contributed equally to the study (Chantal Philippe, Adam Valcek).

## Abstract

We assembled the whole genome sequence (WGS) of a collection of 43 non-redundant modern clinical isolates and four broadly used reference strains of *Acinetobacter baumannii*. Comparison of these isolates and their WGS confirmed the high heterogeneity in capsule loci, sequence types, the presence of virulence and antibiotic resistance genes. However, a significant portion of clinical isolates strongly differ when compared to several reference strains in the light of colony morphology, cellular density, capsule production, natural transformability and *in vivo* virulence. These genetic and phenotypic differences between current circulating strains of *A. baumannii* and established reference strains could hamper the study of *A. baumannii* as an entity. The broadly used reference strains led to the current state of the art of the *A. baumannii* field, however, we propose that established reference strains in the *A. baumannii* field should be carefully used, because of the high genetic and phenotypic heterogeneities. In this study, we generated a collection of high-quality nucleotide sequences of 43 modern clinical isolates with the corresponding multi-level phenotypic characterizations. Beside the contribution of novel fundamental observations generated in this study, the phenotypic and genetic data, along with the bacterial strains themselves, will be further accessible using the first open access online platform called “Acinetobase”. Therefore, a rational choice of modern strains will be possible to select the ones that suit the needs of specific biological questions.

## Introduction

Antibiotics overuse, along with fewer new therapeutics, represents a major challenge for human health (Hernando-Amado et al., 2019; Poirel and Nordmann, 2006). Multidrug-resistant (MDR) bacteria are increasingly isolated around the world, leaving physicians increasingly facing no therapeutic option (Murray et al., 2022). As a direct consequence, patients are currently dying from previously treatable diseases. In this context, WHO and CDC prioritize problematic bacterial pathogens for which antibiotic resistance significantly impacts human health (Centers for Disease Control, 2019; Tacconelli et al., 2018). *Acinetobacter baumannii* (Whiteway et al., 2022), a member of the ESKAPE multidrug-resistant and most problematic nosocomial pathogens (de Oliveira et al., 2020), was designated as a top-priority and critical agent for which therapeutic alternatives are urgently required (Tacconelli et al., 2018).

*A. baumannii* is a Gram-negative opportunistic bacterial pathogen that thrives in hospital settings, especially in intensive care units where weakened patients are treated (Whiteway et al., 2022). Beside their intrinsic and acquired antibiotic resistance, *A. baumannii* resistance to desiccation and disinfectants render any decontamination strategy a real challenge (Chiang et al., 2018). One key aspect sustaining the rapid spread of antibiotic resistance amongst *A. baumannii* isolates is their natural competence (Vesel and Blokesch, 2021). *A. baumannii* bacteria undergo horizontal gene transfer during which exogenous DNA is taken up and integrated within the bacterial genome (da Silva and Domingues, 2016). As a direct consequence, rapidly evolving *A. baumannii* bacteria show a dynamic genome, with an estimated core (conserved) genome of only 16.5%, while 25% of the genome is unique to each strain, having no counterpart in any other *A. baumannii* genome (Imperi et al., 2011). However, despite their clinical relevance, *A. baumannii* bacteria remain poorly understood. Especially, their virulence and non-antibiotic associated resistance still needs to be better characterized (Harding et al., 2017).

One possible explanation for this gap of knowledge is the heterogeneity amongst *A. baumannii* isolates, rendering the multiple approaches of typing of this bacterial species difficult, with as a proof-of-concept the huge diversity of the polysaccharide capsules and the outer core of the lipooligosaccharide (LOS) in *A. baumannii*, in addition to the two multilocus sequence typing (MLST) schemes (Gaiarsa et al., 2019). The LOS is present in *A. baumannii* instead of the common lipopolysaccharide (LPS). Unlike LPS, the LOS is lacking the O-antigen and is composed of lipid A with variable amounts of inner and outer core sugars (Kenyon et al., 2014). The outer sugars of LOS show diversity across strains, dependent on glycosyltransferases and nucleotide-sugar biosynthesis enzymes encoded in a highly variable outer core locus (OCL) (Geisinger et al., 2019). The LOS, a major virulence component in Gram-negative bacteria is encoded chromosomally and to date 14 variants (differing in the presence or absence of genes encoding multiple glycosyltransferases and other enzymes) were described. OCL types can be divided in two groups (A or B) based on the presence of *pda1* and *pda2* genes (Kenyon et al., 2014). The OCL consists of genes involved in the synthesis, assembly and export of complex oligosaccharides that are then linked to lipid A to form the LOS (Kenyon and Hall, 2013). Bacterial capsule consists of a polysaccharide layer deposited as the outermost surface exposed leaflet on prokaryotic cells, impacting the virulence and antibiotics, as well as non-antibiotics-based resistances (Dong et al., 2006; Niu et al., 2019). The locus encoding the genes involved in the production and the assembly of the polysaccharide capsule (KL) is chromosomally encoded and typically ranges from 20 to 35 kb in size. Its genetic organization comprises three modules (Kenyon and Hall, 2013) encoding (i) the export machinery consisting of three proteins; Wza, Wzb, and Wzc, (ii) glycosyltransferases and the capsule processing genes (and optionally genes for the synthesis and modification of complex sugars) and (iii), genes involved in the synthesis of simple sugar substrates.

Witnessing the high diversity amongst *A. baumannii* isolates, our approach aims at investigating to which extend the use of reference strains actually reflects the dynamic nature and intrinsic heterogeneity of these bacteria in general, at the phenotypic and genotypic level. In this study, we assessed the phenotypic and genetic diversity levels of 43 *A. baumannii* modern clinical isolates, and we compared them to established type strains of *A. baumannii* in the field: AB5075, ATCC17978, ATCC19606 and DSM30011, which are widely used in various studies (Bravo et al., 2016; de Silva et al., 2017; Jacobs et al., 2014; Roussin et al., 2019).

## Materials and Methods

### Modern clinical isolates, reference strains and phylogeny inference

A non-redundant collection of 43 modern (not older than 8 years) clinical isolates from the National Reference Center for Antibiotic-Resistant Gram-Negative Bacilli (CHU UCL-Namur) and four established reference strains: ATCC17978-VUB, ATCC19606-VUB, DSM30011- VUB and AB5075-VUB were studied. The reference strains AB5075, ATCC17978 and ATCC19606 are non-MDR (except AB5075) strains of clinical (osteomyelitis, meningitis and urine, respectively) origin (Gallagher et al., 2015; Harding et al., 2017) while DSM30011 is a non-MDR environmental strain obtained from plant microbiota (Repizo et al., 2017). The reference strains were designated with the extension “-VUB” in order to distinguish the strains and their sequences examined within our study. The selected isolates were described in our previous study (Valcek et al., 2021). In order to increase the heterogeneity of the collection, three carbapenem-susceptible isolates (AB21-VUB, AB169-VUB and AB179-VUB) were added and analyzed as well. We sequenced the whole genome of these isolates using shorts reads (Illumina) combined with long reads [Oxford Nanopore Technologies (ONT)] sequencing techniques to generate *de novo* assembled genomes for each of them.

### Short- and long-read sequencing

Genomic DNA extraction and the sequencing library preparation for short-read 2×250 bp paired-end MiSeq (Illumina) sequencing was performed as described before (Valcek et al., 2021). The DNA for long-read MinION (ONT) sequencing was extracted using Genomic-tip 100/G (Qiagen, Hilden, Germany). The long-read sequencing libraries were prepared using 1D Ligation Barcoding Kit (SQK-LSK109 and EXP-NBD104 ONT, Oxford, UK). Samples were QCed using Qubit (dsDNA BR chemistry, Thermo Fisher Scientific), and Fragment Analyzer, Agilent Technologies (using DNF-464 kit). Average size of the fragments was 45-70 kb. Samples were equimolarly pooled and run per 12 in one run which was always 2x reloaded. MinION flowcells had min. 1200 sequencable pores at the start and initial loading was approximately 35 fmol followed by 2 reloads each after 24h into sequencing. The sequencing was performed on MinION Mk1b (ONT) using R9.4.1 (FLO-MIN106) flowcells.

### Sequence data analysis

The long-reads sequences were demultiplexed and basecalled using Guppy v3.2.2 and subsequently were adaptor, quality (Q≤13) and length (5000 bp) trimmed using Porechop v0.2.2 (https://github.com/rrwick/Porechop) and NanoFilt v2.8.0 (de Coster et al., 2018.), respectively. The short reads (BioProject PRJNA734485) were used to polish the long reads employing Ratatosk v0.7.0 (Holley et al., 2021). The corrected reads were then assembled using Flye v2.9 (Kolmogorov et al., 2019) resulting in circular chromosomal contigs of all isolates. The circularity was verified by mapping short reads to corresponding assemblies in order to observer overlapping sequences. The circular chromosomal contigs were polished using long reads via racon v1.4.20 (https://github.com/isovic/racon) and Medaka v1.2.2 (https://github.com/nanoporetech/medaka) and subsequently by quality (Q≤20) and adaptor trimmed (using Trimmomatic (Bolger et al., 2014)) short reads in three rounds of Pilon (Walker et al., 2014) polishing.

### Antimicrobial resistance genotype and phenotype

The antimicrobial susceptibility testing and characterization of its genetic background was performed and described in our previous study (Valcek et al., 2021).

### Natural competence of the isolates

The natural competence of the clinical isolates and reference strains of *A. baumannii* was assessed by starting an overnight bacterial culture from −80°C stock in 5mL of LB (37°C at 160 rpm). The culture was diluted 1:100 in 2mL microtube (10μL of the bacterial culture + 990μL tryptone solution of 5g/L). Then 3μL of the diluted culture was mixed with 3μL of plasmid DNA (100ng/μL) in 1mL microtube. Subsequently, 3μL of the mix of culture and plasmid DNA was transferred to a 2mL microtube containing 1mL tryptone 5g/L agar 2%. A 3μL of diluted bacterial culture without plasmid DNA was used as a negative control.

After 6h incubation at 37°C, 100μL of tryptone solution at 5g/L was added to the microtube with the culture and plasmid DNA and vortexed gently. A 50μL of the suspension was plated on LB agar plates containing apramycin (50μg/mL) to select transformants. The negative control was plated apramycin LB agar plates too. Colonies were counted after overnight incubation at 37°C.

### Hemolytic and protease activities

To assess the potential hemolytic activities of the different *A. baumannii* isolates, we spotted 5 μl of an O/N (overnight) culture of bacteria previously grown in LB medium for 16 hours at 37°C under constant agitation (175 rpm) on 4 different Blood Agar Plates: (i) Columbia Agar with 5% Horse Blood, (ii) Columbia Agar with 5% Sheep Blood, (iii) Trypticase^™^ Soy Agar II with 5% Horse Blood and (iv) Trypticase^™^ Soy Agar II with 5% Sheep blood, all purchased from BD (Becton, Dickinson and Company, Franklin Lakes, NJ). To test for secreted protease activity, we used the same approaches as described above and spotted the 5 μl of bacteria on LB agar plates containing 2% of Skim Milk Powder for microbiology (Sigma-Aldrich/Merck KGaA, Darmstadt, Germany). Plates were incubated at 25°C and monitored for hemolytic and protease activities after 1, 2 and 6 days of incubation.

### Phylogenetic analysis

The maximum-likelihood tree depicting the relatedness of the isolates was constructed from assembled complete genomes using precited open reading frames obtained by Prokka (Seemann, 2014) as an input for the core-genome alignment created using Roary (Page et al., 2015). RAxML (Stamatakis, 2006) was used for calculation of the phylogenetic tree using general time reversible with optimization of substitution rates under GAMMA model of rate heterogeneity method supported by 500 bootstraps. The phylogenetic tree was visualized in iTOL (Letunic and Bork, 2019).

### Genotypic characterization

The resistance and virulence genes were detected using ABRicate (https://github.com/tseemann/abricate) employing ResFinder 4.1 (Bortolaia et al., 2020), VFDB 2022 (Liu et al., 2022) and MEGARes 2.0 (Doster et al., 2020) databases, respectively with 90% threshold for both gene identity and coverage. The typing of capsule-encoding loci (KL) and lipooligosaccharide outer core (OCL) were determined using Kaptive (Wick et al., 2018; Wyres et al., 2020) after manual curation of the corresponding loci by mapping the short reads on anticipated reference sequence of the KL using Geneious R9 (Biomatters, NZ).

### Macrocolony morphology

5 μl of overnight bacterial suspension (~ 1×10^8^ cells) was plated on Columbia Agar with 5% Sheep Blood purchased from BD (Becton, Dickinson and Company, Franklin Lakes, NJ). The plates were incubated non-inverted for 24h at 25°C and subsequently photographed by a Canon^®^ camera.

### Capsule production

1 ml of overnight culture in a 1.5 ml microtube was centrifuged for 2 min at 7000 rcf. The supernatant was removed, and the pellet was resuspended in 1 ml of PBS. Subsequently, 875 μl of PBS resuspended bacteria were mixed with 125 μl of LUDOX^®^ LS colloidal silica (30 wt. % suspension in H_2_O, Merk) (Ardissone et al., 2014; Kon et al., 2020). This mix was then centrifuged for 30 min at 12.000 rcf and immediately photographically recorded.

### Transmission electron microscopy (TEM)

Transmission electron microscopy (TEM) was used for direct capsule visualization by labeling the capsule of 11 *A. baumannii* isolates, 9 modern clinical isolates and 3 reference strains, which ranges from high to low densities. The fixation and staining of the bacteria were performed as described before (Chin et al., 2018). The cupule with the fixed pellet of bacteria (polymerized for 5 hours) was embedded in resin and polymerized (12h at 37°C, 48h at 45°C and 3 days at 60°C). The ~60 nm slides of the resin were marked with acetate uranyl and placed on the electron microscopy grid.

### Virulence in *Galleria mellonella* model of infection

TruLarv research grade larvae of *G. mellonella* (BioSystems Technology) were stored at 15°C no longer than 5 days after arrival and were incubated for 30 min at 4°C prior to injection. Bacteria from an overnight culture were washed with physiological saline (PS) (0.9% NaCl) and diluted to approx. 1×10^7^ CFU/ml. The larvae were injected with 10 μl of PS containing 1×10^5^ CFU/ml of *A. baumannii* in the last left proleg using a 0.3 ml insulin syringe (BD MicroFine). Each of the nine selected strains of *A. baumannii* were injected into 10 larvae and 10 larvae were injected with PS as a negative control. The experiments were carried in duplicates and the survival (assessed by keratinization and mobility) rate was evaluated each day in a period of 5 days.

## Results

### Genotypical characterization of the isolates

We have previously described the sequence types (ST) and the antibiotic resistance profiles of 40 modern carbapenem-resistant clinical isolates of *A. baumannii* bacteria (Valcek et al., 2021). We completed our strain collection by generating the whole genome sequence (WGS) of three carbapenem-sensitive modern isolates (AB21-VUB, AB169-VUB and AB179-VUB) and four broadly used reference strains in the *A. baumannii* field (AB5075-VUB, ATCC19606-VUB, ATCC17978-VUB and DSM30011-VUB), their phylogenetic tree and antimicrobial resistance genes (Supplementary Figure 1).

We further investigate the genetic background of our extended strain collection and generate polished *de novo* assembled genomes by combining Illumina and Oxford Nanopore Technologies sequencing data. Comparison of the whole genome sequences (WGS) of the 43 modern clinical isolates and four reference strains of *A. baumannii*, identified a core genome containing 2009 genes (Figure 1), representing 14.76% of coding sequences (CDS) from the pan-genome of 13611 CDS. The more *A. baumannii* genomes are analyzed, the more unique genes are identified (Figure 2B) while the number of conserved genes decreased. These findings confirm the great variability of *A. baumannii* bacteria, pointing towards a still open pan-genome of *A. baumannii* bacteria that are often changing. Therefore, we still make the current prediction that, every time a new isolate of *A. baumannii* is sequenced, most likely (a) novel gene(s) would be identified.

**Figure 1:**
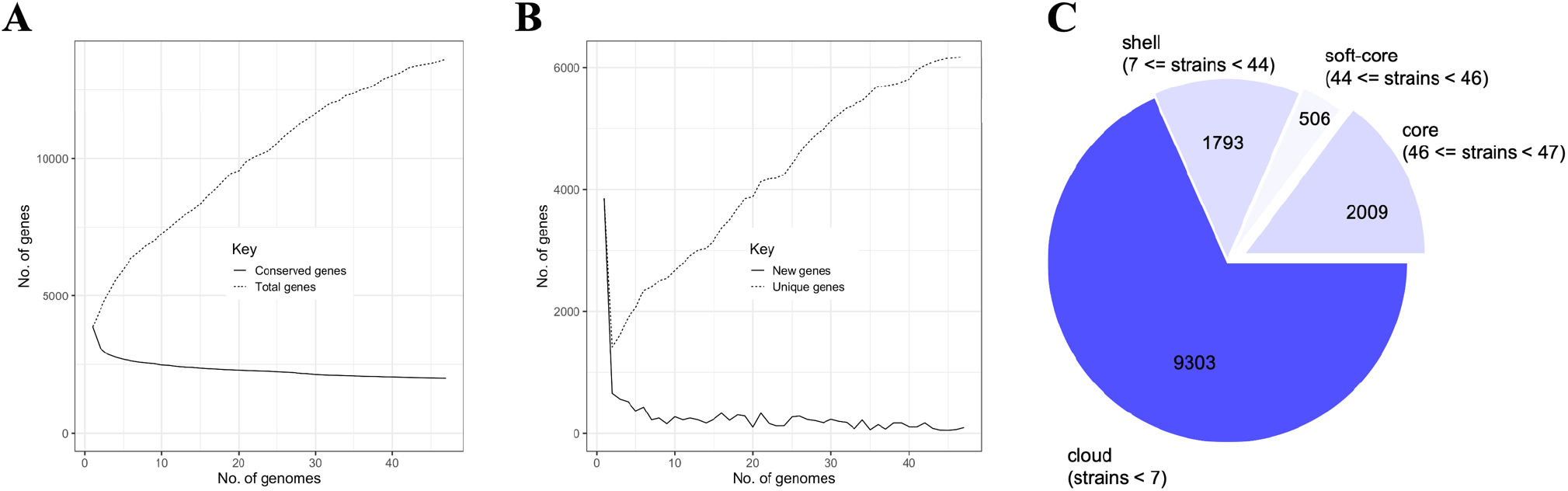
A depiction of (A) Conserved and total gene numbers in the pangenome. (B) Novel and unique gene numbers in the pangenome. These graphs indicate how the pangenome varies as genomes are added. (C) Pangenome pie chart showing the number of core and accessory genes. Accessory genes were divided into soft core, shell and cloud genes.

**Figure 2:**
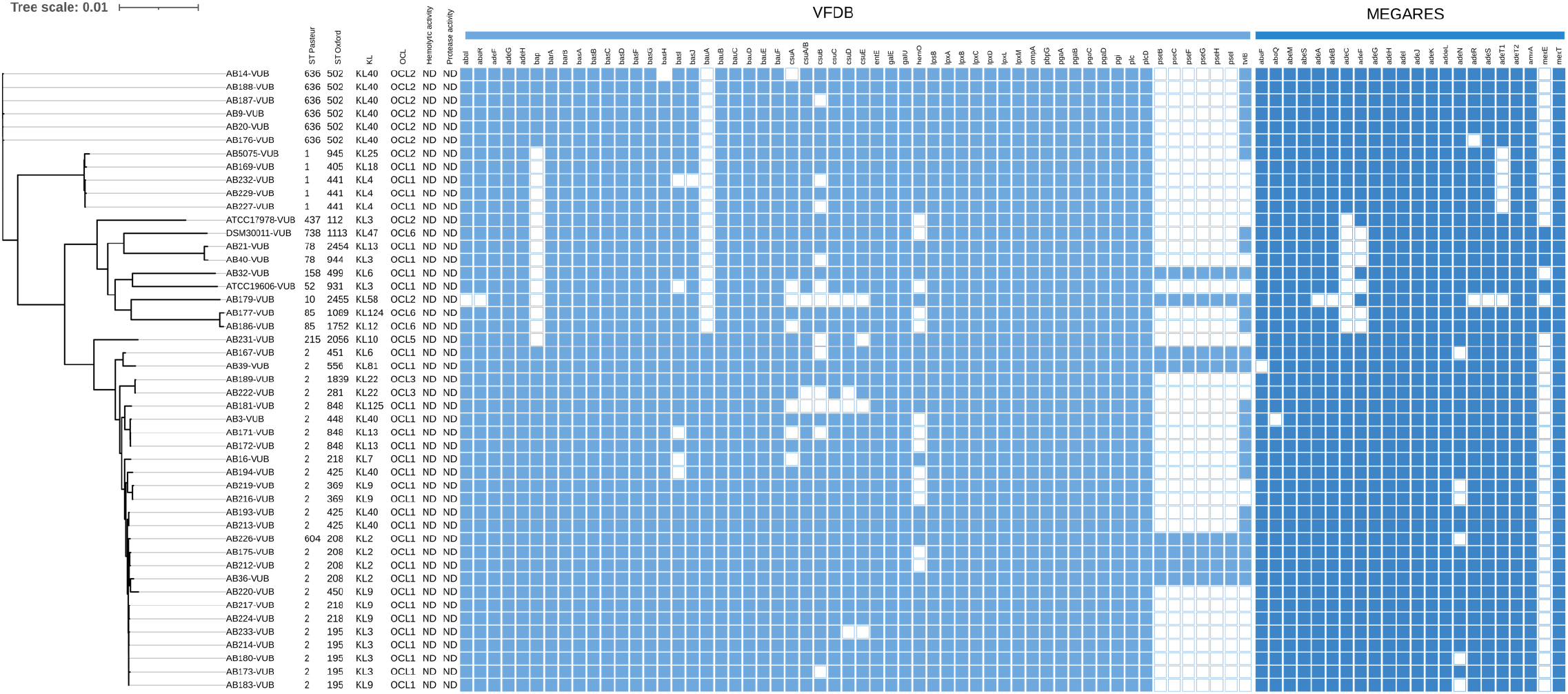
A phylogenetic tree of 43 modern clinical isolates and four reference strains of *A. baumannii* with depiction of ST^Pas^ and ST^Ox^, KL, OCL, hemolytic activity, protease activity and virulence genes as detected using VFDB 2022 and MEGARes 2.0 databases, respectively. ND – Not Detected

We further analyzed the 47 genomes of our strain collection in regard to their respective sequence type (ST), capsule locus type (KL) and lipooligosaccharide locus (OCL) types. Twelve ST groups were identified, with ST2 as the most prevalent (25/43) followed by ST636 (6/43), ST1 (4/43), ST85 (2/43), ST78 (2/43) and ST604, ST215, ST158 and ST10 (one isolate each) (Figure 2). The four reference strains belong to ST1 (AB5075-VUB), ST52 (ATCC19606-VUB), ST437 (ATCC17978-VUB) and ST738 (DSM30011-VUB). We found that the most frequent capsule type was KL40 (10/43) followed by KL9 (6/43), KL3 (5/43), KL2 (4/43), KL13 and KL4 (3/43), KL22 and KL6 (2/42) and KL125, KL124, KL81 KL58, KL18, KL12, KL10 and KL7 (one isolate each) (Figure 2). We confirmed that the reference strains ATCC19606-VUB and ATCC17978-VUB, both belonged to KL3, while DSM30011- VUB to KL47 and AB5075-VUB to KL25 (Arbatsky et al., 2015; Senchenkova et al., 2015; Wang-Lin et al., 2017). In this regard, the reference strains ATCC19606-VUB and ATCC17978-VUB, both KL3, were more representative than DSM30011-VUB (KL47) and AB5075-VUB (KL25) which were sole to be of said KL. The typing of the locus encoding the OCL of the LOS revealed high occurrence of OCL1 (31/43) followed by OCL2 (7/43), OCL6 and OCL3 (2/43) and OCL5 (1/43). We confirmed that the reference strains belonged to OCL2 (AB5075-VUB and ATCC17978-VUB), OCL3 (ATCC19606-VUB) and OCL6 (DSM30011- VUB).

The phylogenetic along with the genetic analyses of KL and OCL showed that, except for AB5075-VUB, the reference strains ATCC17978-VUB, ATCC19606-VUB and DSM30011- VUB only clustered with the clinical isolates of rare ST and KL. Despite AB5075-VUB was sole of the KL25 it clustered closer to the modern clinical isolates of *A. baumannii*. In addition, a variety of genes from operons encoding RND efflux pumps involved in virulence were not present in said three oldest reference strains (ATCC19606-VUB, ATCC17978-VUB and DSM30011-VUB), and a key gene involved in biofilm formation (*bap*) was undetected in all reference strains (Figure 2). Employing MEGARes 2.0 database, only reference strain AB5075- VUB encoded all three RND efflux pumps *adeIJK, adeABC*, and *adeFGH*, while DSM30011- VUB, ATCC17978 and ATCC19606-VUB had an incomplete *adeABC* (lacking *adeC*, all three strains) and/or *adeFGH* (lacking *adeF*, DSM30011-VUB, ATCC19606-VUB) operon. Besides these three reference strains, only six modern clinical MDR strains of *A. baumannii* contained one or more incomplete operons encoding RND efflux pumps.

### Natural competence of *A. baumannii* clinical isolates and strains

We next assessed the natural competence ability of all the *A. baumannii* strains. The reference strain AB5075-VUB was used as a positive control due to its high competence in the tested conditions (Le et al., 2021). We have categorized the strains in regard to its level of competence (Table 1). These results are showing not only variability in the natural competence of the clinical isolates and reference strains of *A. baumannii*, but also a different level of the natural competence.

**Table 1:** A table summarizing number of colonies of *A. baumannii* clinical isolates and reference strains detected after the transformation with plasmid containing apramycin cassette. High: >50 colonies; Medium: 30 – 50 colonies; Low: <5 colonies; Not detected – 0 colonies; in the tested conditions. ND – Not Detected.

| Strain | Level of competence | Strain | Level of competence |
| --- | --- | --- | --- |
| DSM30011-VUB | Medium | AB181-VUB | ND |
| ATCC17978-VUB | ND | AB183-VUB | ND |
| ATCC19606-VUB | ND | AB186-VUB | High |
| AB3-VUB | ND | AB187-VUB | ND |
| AB9-VUB | ND | AB188-VUB | Low |
| AB14-VUB | ND | AB189-VUB | High |
| AB16-VUB | ND | AB193-VUB | ND |
| AB20-VUB | High | AB194-VUB | ND |
| AB21-VUB | ND | AB212-VUB | ND |
| AB32-VUB | Low | AB213-VUB | ND |
| AB36-VUB | ND | AB214-VUB | ND |
| AB39-VUB | ND | AB216-VUB | ND |
| AB40-VUB | ND | AB217-VUB | ND |
| AB167-VUB | ND | AB219-VUB | ND |
| AB169-VUB | High | AB220-VUB | ND |
| AB171-VUB | ND | AB222-VUB | High |
| AB172-VUB | ND | AB224-VUB | ND |
| AB173-VUB | ND | AB226-VUB | ND |
| AB175-VUB | ND | AB227-VUB | High |
| AB176-VUB | High | AB229-VUB | High |
| AB177-VUB | ND | AB231-VUB | ND |
| AB179-VUB | ND | AB232-VUB | High |
| AB180-VUB | ND | AB233-VUB | ND |

### Colony morphology of *A. baumannii* and the need for classification to define new phenotypic categories

We have now confirmed the high heterogeneity at the genetic level amongst our strain collection of MDR modern clinical isolates (Figure 1). To test whether this observation correlates with different levels of phenotypic heterogeneity, we spotted each strain on different blood and milk derived media and observed the potential hemolytic or secreted protease activities (Van der Henst et al., 2018) as well as the ultrastructure of macrocolonies (Figure 3). The clinical isolates and the reference strains did not show detectable hemolytic or protease activities in the tested conditions, which is a common trait amongst *A. baumannii* bacteria (Bouvet and Grimont, 1986) (Figure 1). The reference strains AB5075-VUB, DSM30011- VUB, ATCC17978-VUB and ATCC19606-VUB do represent this common phenotypic trait, despite the high genetic diversity observed, confirming that the tested modern clinical isolates did not acquire increased hemolytic nor secreted protease activities. However, the analysis of the macrocolonies morphology highlights an important diversity level of the modern clinical isolates compared to the reference strains. As shown in Figure 3 and Table 2, we observed at least five different categories of ultrastructure of macrocolony for the modern clinical isolates and only two categories for the tested reference strains, with the more modern isolate AB5075- VUB being different compared to the three other reference strains DSM30011-VUB, ATCC17978-VUB and ATCC19606-VUB. We could identify six constitutive mucoid strains in the macrocolony type (MT) “A” group of *A. baumannii* isolates (Figure 3B). The colonies of the isolates with constitutive mucoid phenotype had circular shape and smooth margin. The least frequent morphology of a colony and second most mucoid was circular with raised top of darker white color with only three isolates, representing the MTB group (Figure 3B). The most abundant (n=17) type of colony (MTC) was irregular with translucent bottom and opaque top with irregular margin. This group also includes the most modern reference strain AB5075-VUB (Figure 3A). The isolates of the MTD group (n=14) produced circular colonies with undulate margin and volcano shaped center. Most of the reference strains belongs to the MTE group of seven least mucoid isolates (Figure 3B) with circular shape, translucent center and opaque outer ring. Hence, the shapes of the colonies allowed to divide the isolates into five macrocolony type groups with one representative isolate depicted in Figure 3 (Table 2).

**Figure 3:**
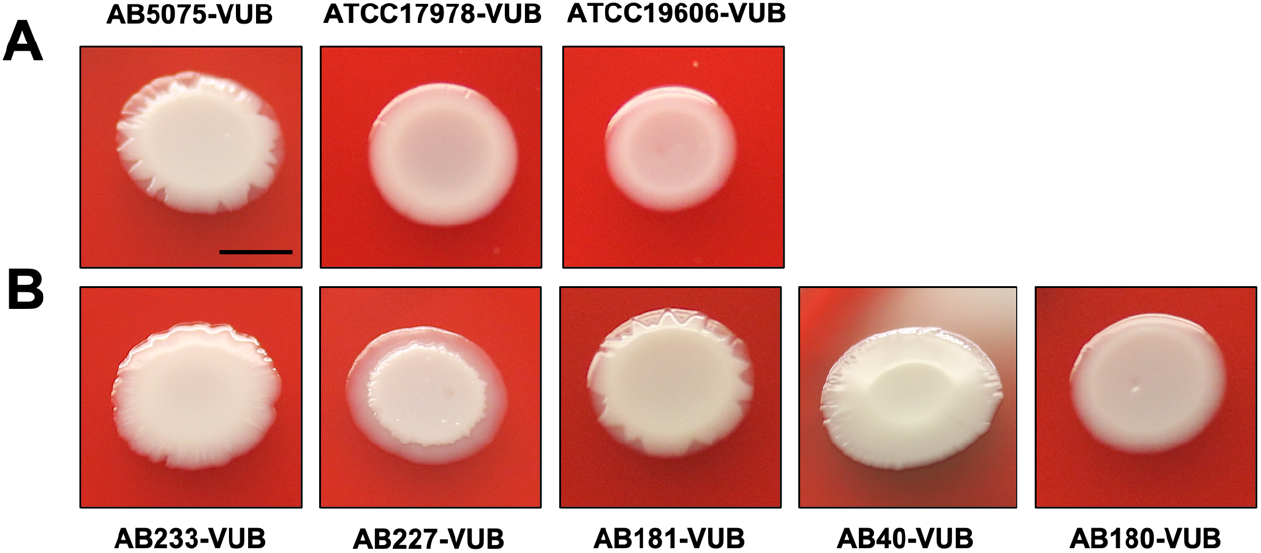
Diversity of the macrocolony types of the reference strains (A) compared to modern clinical isolates (B) grown on columbia agar plates with 5% sheep blood. Scale bar: 1 cm.

**Table 2:** A table dividing the reference strains and the clinical isolates to five categories (MTA-MTE) based on the colony morphology from the most mucoid phenotype (A) to the least mucoid phenotype (E). Each category is represented by one isolate, which is depicted in Figure 3B.

| Macrocolony Type (MT) | AB233-VUB (MTA) | AB227-VUB (MTB) | AB181-VUB (MTC) | AB40-VUB (MTD) | AB180-VUB (MTE) |
| --- | --- | --- | --- | --- | --- |
|  | AB3-VUB | AB16-VUB | AB14-VUB | AB9-VUB | AB32-VUB |
|  | AB39-VUB | AB20-VUB | AB21-VUB | AB36-VUB | AB212-VUB |
|  | AB173-VUB |  | AB169-VUB | AB167-VUB | AB213-VUB |
|  | AB177-VUB |  | AB171-VUB | AB175-VUB | ATCC17978-VUB |
|  |  |  | AB172-VUB | AB176-VUB | ATCC19606-VUB |
|  |  |  | AB186-VUB | AB179-VUB | DMS30011-VUB |
|  |  |  | AB188-VUB | AB183-VUB |  |
|  |  |  | AB193-VUB | AB187-VUB |  |
|  |  |  | AB194-VUB | AB189-VUB |  |
|  |  |  | AB217-VUB | AB214-VUB |  |
|  |  |  | AB219-VUB | AB216-VUB |  |
|  |  |  | AB220-VUB | AB222-VUB |  |
|  |  |  | AB224-VUB | AB226-VUB |  |
|  |  |  | AB229-VUB |  |  |
|  |  |  | AB231-VUB |  |  |
|  |  |  | AB232-VUB |  |  |
|  |  |  | AB5075-VUB |  |  |

### Capsule production of *A. baumannii*

As the mucoid phenotype can reflect the presence of an abundant polysaccharide capsule surrounding bacteria, we established a density gradient assay to assess encapsulation level of all *A. baumannii* isolates in a medium to high throughput level (Figure 4) (Whiteway et al., 2021). In this phenotypic assay, low density bacteria have a high capsulation level, while denser bacteria have lower capsulation levels (Figure 4A). A high heterogeneity degree concerning the density of *A. baumannii* bacteria, crossing the full range from low to high density levels is observed (Figure 4D). Interestingly, four clinical isolates were divided into two fractions, suggesting diversity in production of CPS even within the same isolate, possibly pointing towards a high frequency of phase variation leading to phenotypic heterogeneity previously described (Tipton et al., 2015). In majority, the reference strains show high densities except for AB5075-VUB characterized by a medium density level. Despite having low capsulation levels, the ATCC19606-VUB isolate constantly shows a lower density compared to the strain ATCC17978-VUB. On the other hand, none of the high-density isolates belonged to KL40, unlike majority of low-density producers, pointing towards causality. While in high density isolates no other pattern than frequent occurrence of ST2 (10/15) was observed, the isolates of low density were often of KL40 (7/9) of ST2 (5/9) or ST636 (4/9). In conclusion, we could not attribute a specific cell density with the capsule type. The colony morphology (MTE) and the density gradient of four modern clinical isolates of our collection (AB32-VUB, AB180-VUB, AB212-VUB and AB213-VUB) resembles those of the low capsulated reference strains ATCC17978-VUB, ATCC19606-VUB and DSM30011-VUB, hence was to a certain extent represented by two reference strains. In this regard, the reference strain AB5075-VUB represented 17 clinical isolates of MTC, yet the density gradients of this group were too heterogenous to be represented by a sole reference strain AB5075-VUB. However, the majority (n=22/43) of the modern clinical isolates was not represented by the reference strains of *A. baumannii* considering the capsule production (density gradient) and macrocolony types.

**Figure 4:**
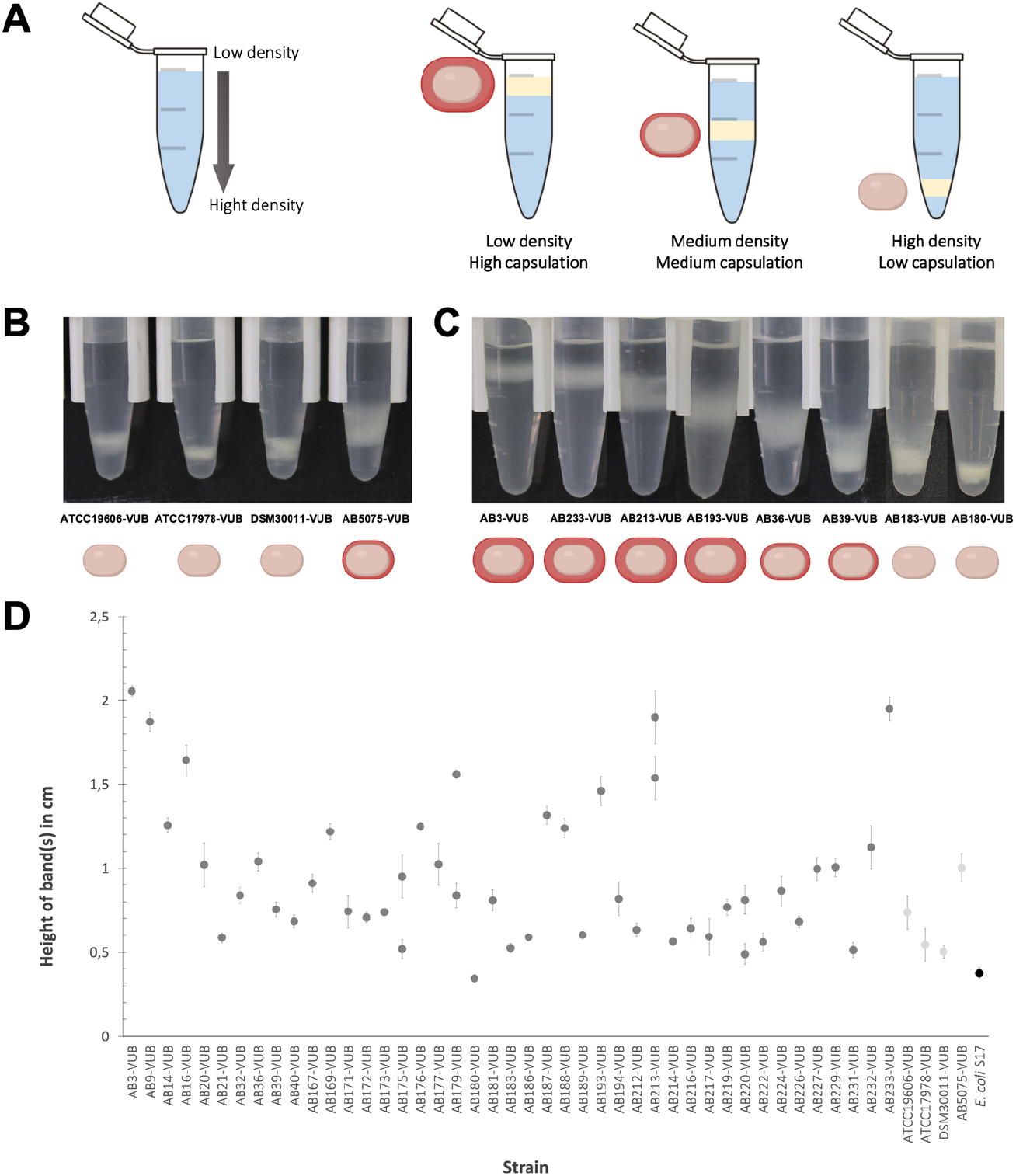
(A) Depiction of the density gradient measuring principle. (B) Capsulation level of the reference strains and (C) comparison of various levels of capsulation of the modern clinical strains of *A. baumannii*. (D) Quantification of the density of bacterial cells measured in a gradient colloidal silica; standard deviation was calculated from biological triplicates.

To directly observe capsule deposition and thickness at high resolution and at the single cell level, we combined capsule labeling with transmission electron microscopy (TEM) approach (Figure 5). As expected, high density bacteria are surrounded by a denser and thicker capsule compared to low density bacteria (Figure 5B). Concerning the reference strains, we confirm AB5075-VUB to be the more capsulated reference strain tested in this study. We could not detect any capsule deposition on the ATCC17978-VUB strain and the ATCC19606-VUB shows weak and heterogeneously deposited layer in the tested conditions. This is in line with the reference strain ATCC19606-VUB being less dense compared to the reference strain ATCC17978-VUB in our density gradient assay (Figure 4D). Two less capsulated modern strains, the AB180-VUB and AB183-VUB also show high densities (in density gradient assay) with less abundant capsule deposition. The least dense isolates AB3-VUB, AB193-VUB and AB213-VUB show a high abundance of capsule formation. Accordingly, the strains AB36- VUB that shows an intermediate density level have an intermediate capsule deposition level while the AB39-VUB strain reaches a higher density with less capsule deposition. Taken together, these data show that bacterial density correlates with capsule abundance of *A. baumannii* bacteria and that the reference strains does not show all the heterogeneity observed in the modern clinical isolates tested regarding capsule production level.

**Figure 5:**
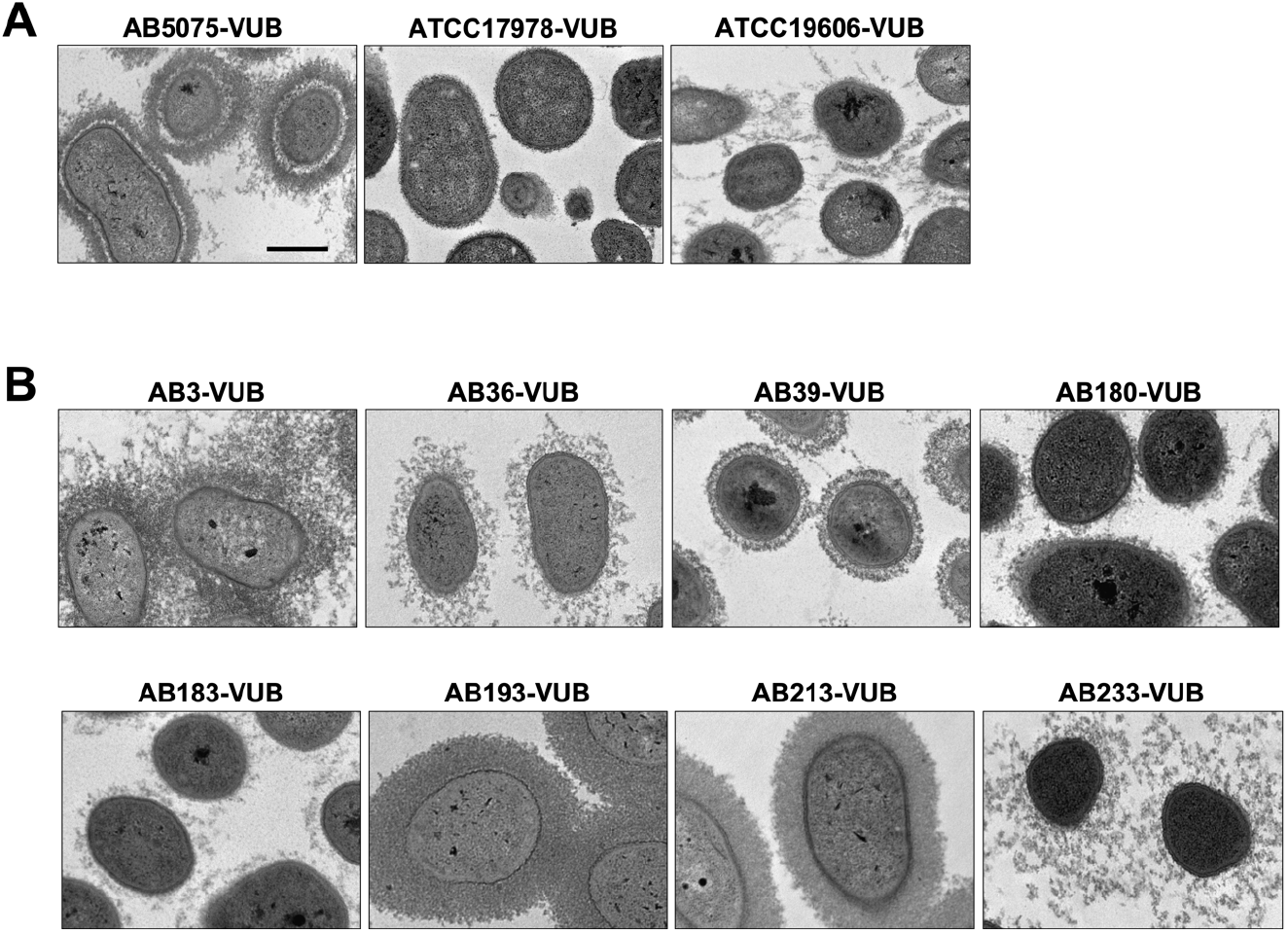
A TEM direct capsule visualization of three reference strains (A) and 8 modern clinical strains (B) of various cell densities. Scale bar: 500 nm.

### Constitutive mucoid strains are not hyper-virulent

As constitutive mucoid strains were observed only amongst the group of the modern MDR clinical isolates, we wondered if this phenotype could be associated with a higher *in vivo* virulence potential. *Galleria mellonella* larvae were infected with highly, medium, and weakly capsulated isolates, while the AB5075-VUB reference strain was used as a positive control of virulence (Jacobs et al., 2014). Concerning the constitutive mucoid isolates, while the AB213- VUB is highly virulent, the constitutive mucoid isolates AB3-VUB and AB233-VUB are weakly virulent (Figure 6). The medium capsulated isolate AB39-VUB was highly virulent and killed 100% of the larvae within one day, being even more virulent than the AB5075-VUB, while the medium capsulated AB36-VUB showed an intermediary virulence level. Interestingly, the low capsulated isolates (AB183-VUB and AB180-VUB) were the less virulent, AB180-VUB being avirulent in the tested conditions (Figure 6). These data show that a constitutive high capsulation and mucoid phenotype does not correlate with a higher virulence potential in *A. baumannii*. However, this highlights that low capsulation levels characterizes weakly virulent phenotypes, showing that *A. baumannii* bacteria required capsule deposition for full virulence. This is in agreement with previous studies showing that genetically manipulated capsule deficient *A. baumannii* strains have decreased virulence levels (Talyansky et al., 2021; Whiteway et al., 2021). As the virulence does not directly correlate with a specific KL or OCL type, we can conclude that the KL or OCL are not solely responsible for *A. baumannii* virulence in the tested conditions, which might require multifactorial influence. Taken together, these observations show that the reference strain AB5075 is a good bacterial model to study the virulence potential in *G. mellonella* while the less virulent strains ATCC19606 and ATCC17978 may be used in studies tackling down the modern clinical isolates exhibiting decreased virulence in *G. mellonella*.

**Figure 6:**
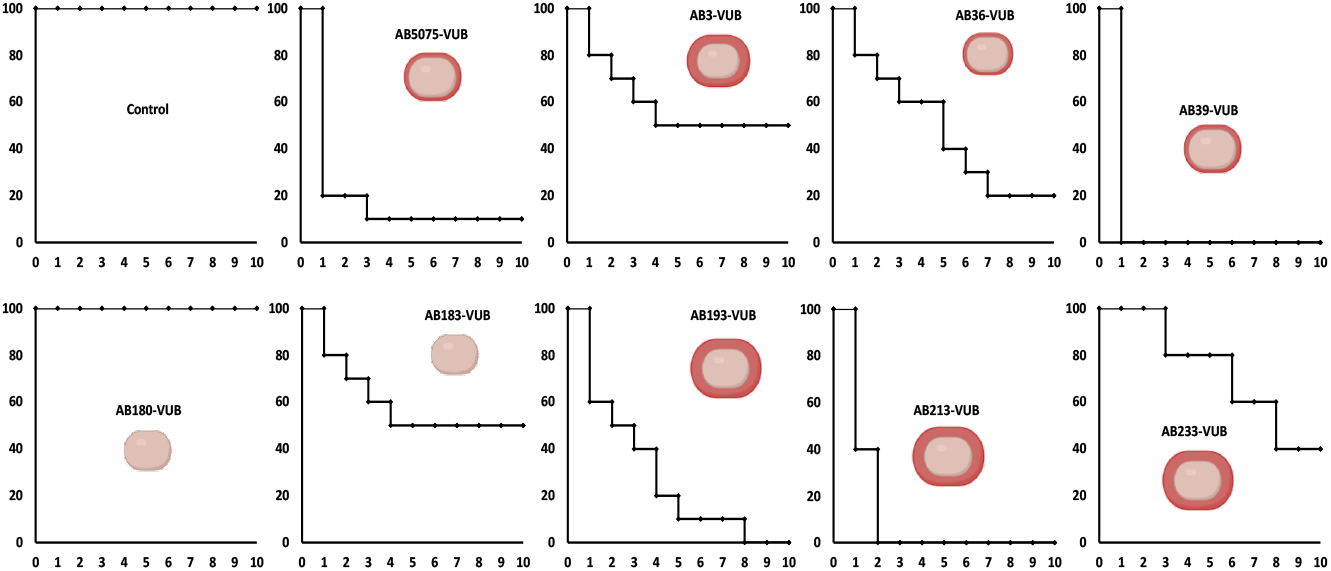
*In vivo* virulence of several modern *A. baumannii* isolates determined in *G. mellonella* model. X axis, days after inoculation and Y axis, percentage of living *G. mellonella*. The control condition is the PS without bacteria.

## Discussion

The 43 modern MDR clinical isolates and four reference strains of *A. baumannii* were whole-genome sequenced, *de novo* assembled and analyzed in the context of genotype and phenotype. We identified 12 different ST groups in our collection of 43 modern clinical isolates. While the reference strain AB5075-VUB belonged to less frequently occurring ST1 the reference strains ATCC19606-VUB (ST52), ATCC17978-VUB (ST437) and DSM30011-VUB (ST738) did not reflect the current trends of ST in clinical settings but rather rare sequence types. The lack of representation of the ST of the clinical isolates is expectable only in the case of DSM30011- VUB as this strain has environmental origin (plant microbiota). The ST2 group, previously described as one important and clinically relevant group is also the most widely disseminated ST among the complete and draft genomes available (Hamidian and Nigro, 2019). Most of the modern clinical isolates of our collection belonged to ST2, yet none of the reference strains represented this sequence type.

Our phylogenetic analysis (Figure 1) identified several clades, mostly in accordance with the ST of the isolates. However, the reference strains DSM30011-VUB, ATCC17978-VUB and ATCC19606-VUB formed a cluster with the least abundant sequence types of the clinical isolates of *A. baumannii* [except AB226-VUB (ST604)]. Only this cluster consisted of the isolates which all lacked *adeC* and *bauA* genes which are discussed below, suggesting an extreme genetic diversity which represents only a minor proportion of the current clinical isolates of *A. baumannii*. On the other hand, from the four reference strains, only AB5075-VUB was the most similar to the modern clinical strains of *A. baumannii* in regard to OCL and virulence genes. When considering AB5075-VUB capsule production, its phenotype was resembling phenotypes of the clinical isolates with low production. Nevertheless, similarly to the other reference strains, it lacked *bauA* and *adeC*. The lack of the combination of these two virulence genes was also detected in ten modern clinical isolates proving that AB5075-VUB still shares partial similarity with circulating clinical strains of *A. baumannii*. The described and listed virulence genes in our study only account for a small amount of the virulence genes as only known can be detected and possibly novel virulence genes remain to be identified. Noteworthy, regardless of observed discrepancy in detection of *adeFGH* operon using VFDB 2022 and MEGARes 2.0 databases, the absence of *adeFGH* was described previously (Ardehali et al., 2019; Coyne et al., 2010).

Capsule heterogeneity is a landmark of *A. baumannii* bacteria. The surface polysaccharides play key roles in fitness and virulence of *A. baumannii* and protects it from the environment, increases resistance to antimicrobial compounds and helps to evade the host immune system (Geisinger and Isberg, 2015; Weber et al., 2016). There are more than 137 KL types identified so far (Kenyon and Hall, 2021), and this variability demonstrates the diversity which must be surveilled in the modern clinical isolates as the capsular polysaccharide is a potential target for therapeutical agents and vaccines (Yang et al., 2017). In our collection of 43 modern clinical isolates of MDR *A. baumannii*, KL40 (10/43) dominated while KL3 was detected in five isolates.

In our study, the most prevalent locus encoding outer-core lipooligosaccharide (OCL) type in the modern clinical isolates of MDR *A. baumannii* was OCL1 (31/43). However, none of the reference strains represented this most prevalent OCL1 type as they belonged to OCL2 (AB5075-VUB and ATCC17978-VUB), OCL3 (ATCC19606-VUB) and OCL6 (DSM30011- VUB) suggesting their rarefaction in following the current epidemiological trends.

Two reference strains (DSM30011-VUB and AB5075-VUB) in this study lacked *bap* gene encoding biofilm associated protein which could influence the studies exploring biofilm properties using these reference strains. However, the gene *bap* was found in ATCC17978- VUB and ATCC19606-VUB with 86% identity (VFDB 2022) same as in three clinical isolates (AB40-VUB, AB32-VUB and AB21-VUB), therefore assigned as undetected in the Figure 1 as the identity falls under the applied threshold if 90%. The *bap* encodes protein required for formation of three-dimensional biofilm towers and water channels on abiotic and biotic surfaces such as polypropylene, polystyrene, and titanium (Brossard and Campagnari, 2012). Bap protein is also involved in adherence of *A. baumannii* to human bronchial epithelial cells and human neonatal keratinocytes (Brossard and Campagnari, 2012). Moreover, nearly half of the examined strains (20/47) including four reference strain did not carry *bauA* encoding iron-regulated outer membrane protein BauA. BauA protein provides protection against sepsis caused by *A. baumannii* and was also identified as a vaccine candidate (Aghajani et al., 2019; Ni et al., 2017). Interestingly, six strains (including two reference strains DSM30011-VUB, ATCC19606-VUB and four clinical isolates AB177-VUB, AB186-VUB, AB40-VUB and AB21-VUB) encoded membrane fusion protein MexE protein, part of RND efflux pump from *Pseudomonas aeruginosa* (Köhler et al., 1999). Notably, the very same six strains were lacking a gene encoding membrane fusion protein AdeF from AdeFGH RND efflux pump, possibly restoring its function as AdeF and MexE share 50% homology. However, using the VFDB 2022 database, all clinical isolates and reference strains encoded *adeFGH* operon.

The deficiency of the clinical isolates and the reference strains in hemolytic and protease activities is in agreement with the fact that only some species of *Acinetobacter* genus, e.g., *Acinetobacter haemolyticus* (Touchon et al., 2014) are capable of such phenotype. Even despite this shared phenotypical trait, there was a high level of diversity of macrocolonies, where the reference strains represented only two variants out of five types observed in the clinical strains. Several modern isolates show a constitutive mucoid phenotype that is not observed in the reference strains. Isolates belonging to the same clusters (Figure 1) do not show the same ultrastructure, showing a high diversity even within the same phylogenetic group. To be noted, also external events such as gene disruption can affect the phenotype of the macrocolony, as was observed in AB5075-UW by Perez-Varela (Pérez-Varela et al., 2020) by disrupting *relA* ortholog (*ABUW_3302*) using transposon insertion. Taken together, this points out the insufficiency of the reference strains in grasping the complete phenotypical (and genotypical) diversity of the modern clinical isolates.

The natural competence for transformation is one of the ways of horizontal gene transfer *A. baumannii* uses to acquire extracellular DNA from the environment and incorporates it into its own genome via homologous recombination (Dubnau and Blokesch, 2019). We observed that the competence to be naturally transformed varied within clinical isolates of *A. baumannii* from which 13 were transformed, 18 were not and 12 were intrinsically resistant to apramycin, therefore unable to assess. This diverse trend was copied by the reference strains as well as AB5075-VUB and DSM30011-VUB were successfully transformed while ATCC17978-VUB was not and ATCC19696-VUB showed intrinsic resistance to apramycin. There are multiple factors influencing the ability of *A. baumannii* to be naturally transformed such as presence of the H-NS (Le et al., 2021) which was present in each case within our isolates and strains. However, the major role is played by type IV pilus genes (Leong et al., 2017) which are growth phase dependent (Vesel and Blokesch, 2021), and might be the explanation of some isolates and strains not being transformed. However, the tested conditions of natural competence were standardized pointing out further physiological diversity within *A. baumannii* clinical isolates and reference strains. Despite the proven relevance of the clinical isolates of our collection (collected as problematic and modern nosocomial isolates), we cannot rule out a bias to certain extend.

We observe a high diversity in production of the capsular polysaccharide using direct and indirect visualization methods. We show a correlation between capsule abundance and density levels, but correlation does not mean causation. We cannot rule out the possibility that the capsule type *per se* (and not only the capsule abundance) or other factors, influence the density of *A. baumannii* bacteria. The fact that widely used reference strains show low to medium encapsulation degrees add an additional reason why reference strains should be carefully used and argue in favor of considering the strain AB5075-VUB as the best representative bacterial model out of the four reference strains tested in this study. However, a rather complex solution to selection of proper strain to be used in a specific study is needed such as Acinetobase (Valcek et al., 2022). Acinetobase is a comprehensive database providing the community with genotype, phenotype and the strain of *A. baumannii* itself.

The *Galleria mellonella* infection model proved itself as a valuable source of information on the virulence level of *A. baumannii*. This model also highlighted AB5075-VUB to be more virulent than ATCC19606-VUB and ATCC17978-VUB in the tested conditions (Jacobs et al., 2014). These results support the usage of modern clinical isolates for study of virulence, hence other highly virulent isolates such as hypervirulent *A. baumannii* LAC-4 (Ou et al., 2015) should be considered as well.

As constitutive mucoid strains were observed only amongst the group of the modern MDR clinical isolates, we wondered if this phenotype could be associated with a higher *in vivo* virulence potential. The virulence in *G. mellonella* model varied for the tested clinical isolates and the reference strains and while a constitutive mucoid phenotype does not correlate with a higher virulence, we confirm using modern clinical isolates that low capsule production impedes full virulence in *A. baumannii*. These observations do not confirm the results of Shan et al., (Shan et al., 2021) who concluded that the mucoid *A. baumannii* isolates were more hypervirulent than the nonmucoid strains. This discrepancy may point towards multifactorial background of hypervirulence.

The signal(s) or conditions regulating capsule production remain to be determined. Noteworthy, the isolates AB193-VUB and AB213-VUB with higher integrity of the CPS layer (Figure 5) have shown higher virulence in *G. mellonella* (Figure 6) than isolates with dispersed CPS.

The study of genetic features which are not shared by the majority of *A. baumannii* isolates (soft-core, shell and cloud genes (Cummins et al., 2022)) may lead to novel discoveries as well. However, the global impact of the clinical importance (drug therapies or target-driven drug discoveries) will suffer from linking the specific genes and features only to certain isolate or lineage.

## Conclusion

*A. baumannii* are heterogenous bacteria, both at the genetic and phenotypic levels. In this study, we characterized 43 modern clinical isolates from different phylogenetic groups and 4 reference strains with common but also very different behaviors. Therefore, the reference strains tested in our study do not cover the whole heterogeneity found in the modern isolates of *A. baumannii*.The studies previously published using these reference strains built a strong state of the art in the *A. baumannii* field and beyond, showing their usefulness. However, the data presented in our study show that the specific use of one or only a limited subset of reference strains can hinder important processes characterizing clinically relevant isolates and the *A. baumannii* bacteria as a whole. As an answer to that identified pitfall, we propose a variable collection of modern clinical isolates that are characterized at the genetic and phenotypic levels, covering the full range of the phenotypic spectrum, with five different macrocolony type groups, from avirulent to hyper virulent phenotype, and with non-capsulated to hyper mucoid strains, with intermediate phenotypes as well. This will allow selecting a reference strain rationally, facilitated by the new Acinetobase (Valcek et al., 2022) platform, which suits the needs of an ongoing study with a particular biological question. This is especially important for new antimicrobial screening purposes, for which conserved targets amongst a significant proportion of problematic *A. baumannii* isolates is a prerequisite. While strain specific observation remains interesting *per se*, in the context of such a drastic heterogeneity, any new identified target, antimicrobial compound, or fundamental observation deserved to be tested on diverse relevant *A. baumannii* isolates.

## Data availability

The long- and short-read sequences were deposited in GenBank under BioProject PRJNA701627, PRJNA798866 and PRJNA734485, respectively.

## Acknowledgements

We are grateful to the URBM and URBE research groups, as well as the Electron Microscopy Service from UNamur, for the access to their equipment and their expertise. We thank Ivan Mateus from the Department of Ecology and Evolution, University of Lausanne, Lausanne, Switzerland for a fruitful discussion of initial bioinformatical analyses. We thank the National Reference Laboratory for Monitoring of Antimicrobial Resistance in Gram-negative Bacteria, CHU Mont-Godinne, Université Catholique de Louvain (UCL), Yvoir (Belgium) for providing the modern clinical isolates.

## Contribution

CP, CVDH and ER performed phenotypical experiments. AV, CVDH and KN performed bioinformatical analyses. CW extracted DNA for the long-read sequencing. MB, CP, CVDH and TC performed the infections of *Galleria mellonella*. TDP, WDC and MS set up the LRS strategy, sample, library preparation and sequencing. JK provided her expertise in KL and OCL identification. AV and CVDH wrote the manuscript.

---

**Supplementary Figure 1:**
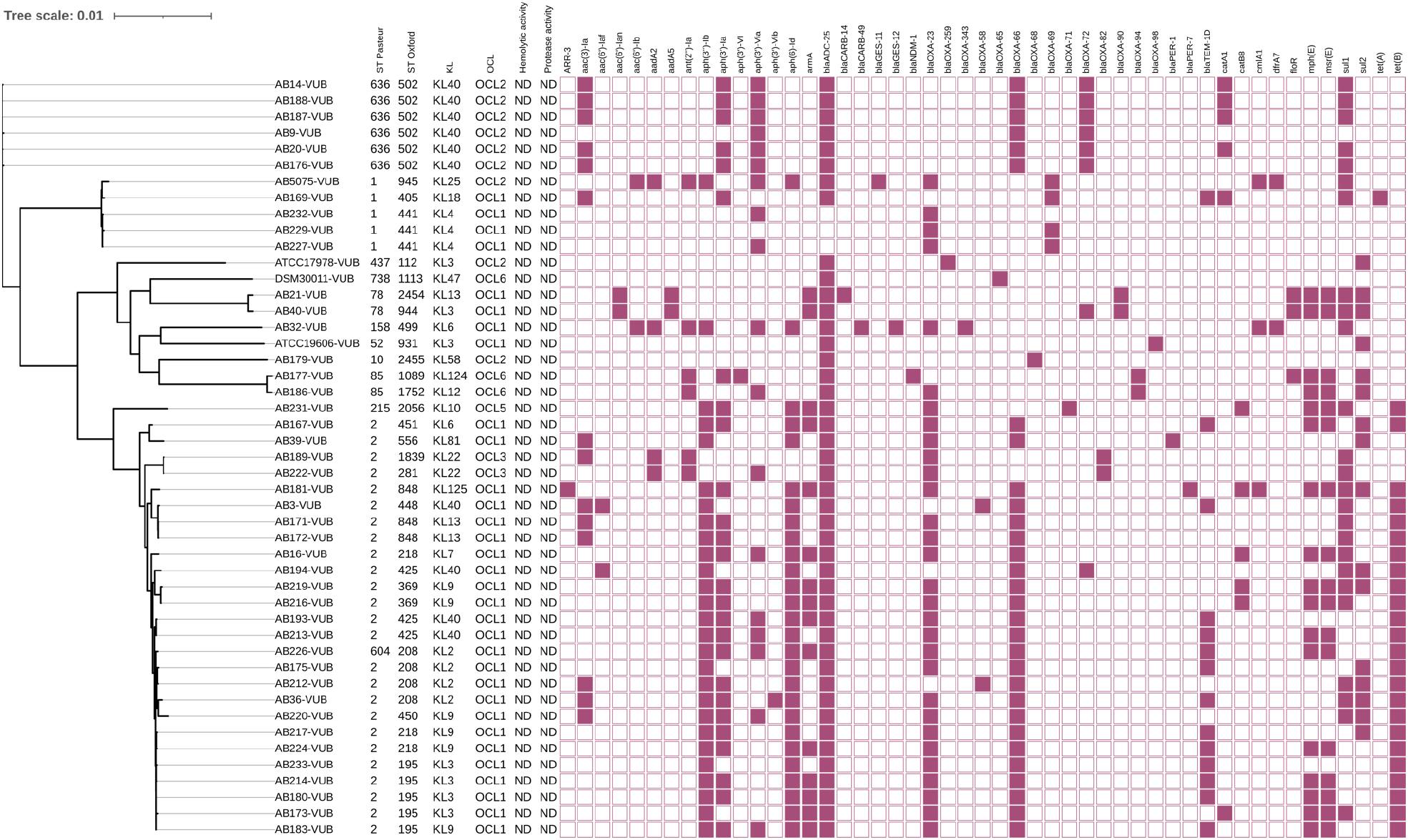
A phylogenetic tree of 43 modern clinical isolates and four reference strains of *A. baumannii* with depiction of ST^Pas^ and ST^Ox^, KL, OCL, hemolytic activity, protease activity and resistance genes, respectively. ND – Not Detected

